# Cortical suppressive waves shape the representation of long-range apparent motion

**DOI:** 10.1101/372763

**Authors:** S Chemla, A Reynaud, M di Volo, Y Zerlaut, L Perrinet, A Destexhe, F Chavane

## Abstract

How does the brain link visual stimuli across space and time? Visual illusions provide an experimental paradigm to study these processes. When two stationary dots are flashed in close spatial and temporal succession, human observers experience a percept of motion. Large spatio-temporal separation challenges the visual system to keep track of object identity along the apparent motion path. Here, we utilize voltage-sensitive dye imaging in primary visual cortex (V1) of the awake monkey to investigate whether intra-cortical connections within V1 can shape cortical dynamics to represent the illusory motion. We find that the arrival of the second stimulus in V1 creates a suppressive wave traveling toward the retinotopic representation of the first. Computational approaches show that this suppressive wave can be explained by recurrent gain control fed by the intra-cortical network and contributes to precisely encode the expected motion velocity. We suggest that non-linear intra-cortical dynamics preformat population responses in V1 for optimal read-out by downstream areas.

## Introduction

When two stationary stimuli are successively flashed in spatially separated positions, it generates the so-called “apparent motion” illusion (Wertheimer 1912). This illusion, well characterized in psychophysics (Burr and Thompson 2011), depends on the spatio-temporal characteristics of the stimulus, being called “short-range” vs “long-range” apparent motion (lrAM) for spatial separation below or above 0.25° and temporal separation below or above 80 ms respectively (Braddick 1980). In psychophysics, intrinsic differences were reported between these two types of apparent motion, however, there is some debate whether it is underlined by same or different process (Cavanagh and Mather 1989). In physiology, while we have a clear idea on the neuronal processing generating direction-selective neuronal response to short-range apparent motion stimuli (Mikami, Newsome, and Wurtz 1986b), we still have a poor understanding of how the visual system process lrAM. This is probably because the spatial separation between individual strokes of the lrAM extend beyond the typical extent of receptive fields in the early visual system, at least in primates. In the case of the lrAM, psychophysicists have long highlighted the necessity to have a process, such as the “reviewing process” (Kahneman, Treisman, and Gibbs 1992), that will link the transient apparitions of stimuli in different spatial and temporal positions in order to generate a coherent motion percept of a single object, hereby solving the problem of “phenomenal identity” (Ternus 1926) or “correspondence” (Ullman 1978). Downstream areas with large receptive fields are a natural expected integration unit for such extended spatiotemporal input. Indeed, it has been recently shown in human that the feedback from MT to V1 plays an important role in the processing of lrAM (Wibral et al. 2009; Muckli et al. 2002; Vetter, Grosbras, and Muckli 2015), as well as evidences of downstream activation along the ventral stream (Zhuo et al. 2003). However, it is still unclear whether and how the “reviewing” process, needed to keep track of the object identity along the motion trajectory, can be achieved within these receptive fields.

As suggested from fMRI experiments in human, the population activity within V1 could participate in formatting the representation of lrAM (Muckli et al. 2005). The extended precise retinotopic map in V1 makes it indeed an ideal platform for representing and creating, at the level of the neuronal population, the trajectory of the apparent motion illusion, a representation that could be read-out by downstream areas (Mumford 1991; Lee et al. 1998). In particular, V1 has the highest resolution (Lee et al. 1998) to achieve the interactions in space and time needed to link the individual strokes of the apparent motion (Lee et al. 1998; Adelson and Bergen 1985). In such context, intra-cortical and inter-cortical connectivity would be the natural substrate to underlie the necessary spatio-temporal interactions (Deco and Roland 2010; Muller et al. 2018). Importantly, these two networks have intrinsically different spatio-temporal properties, the inter-cortical network operating over very large extent but with poor spatial and temporal resolution (Angelucci et al. 2002; Stetter 2002), and the intra-cortical network has a more limited extent but with high spatial and temporal resolution (Muller et al. 2014; Bringuier et al. 1999; Bullier 2001). Furthermore, they constitute the vast majority of synaptic contacts in the cortex, the feedback accounting for less than 20% and the intra-cortical connectivity contributing to 80% of the number of neuronal contacts, while the feedforward less than 1% (Markov et al. 2011). Such connectivity seems therefore like a good candidate to link transient spatio-temporal events (Muller et al. 2018). It was indeed shown, in the anesthetized cat, to shape visual information for a dynamic representations of sequences of static stimuli (Jancke et al. 2004; Gerard-Mercier et al. 2016) likely implicating non-linear gain control on the feedforward input (Reynaud, Masson, and Chavane 2012). However, it is still unclear whether and how the cortico-cortical interactions could participate to shape the representation of lrAM within V1 retinotopic map in the awake monkey.

To answer this question, we used optical imaging of voltage-sensitive dyes (VSDI) in the awake fixating monkey, to measure how V1 neuronal population integrates a two-stroke lrAM that overreached individual neuronal receptive field size. In response to a single stroke, activity in V1 propagates in space and time (Grinvald et al. 1994; Slovin et al. 2002; Sato, Nauhaus, and Carandini 2012; Bringuier et al. 1999; Muller et al. 2014), with spatial and temporal constants that cover about 3 mm and 80 ms. In response to the lrAM of various spatio-temporal separations, we observed the emergence of a direction-selective representation of the lrAM in V1. This representation is the result of a systematic wave of suppression propagating in the opposite direction of the lrAM: initiated at the second stimulus onset and propagating to suppress the residual response to the first stimulus. A computational model was developed to understand the origin of such suppressive waves. It shows that intra-cortical interactions can lead to the observed suppressive waves when two very plausible conditions are met: (i) the inhibitory cells have a higher gain than excitatory cells, and (ii) there is a shunting effect of the associated synaptic conductances. Using a spatio-temporal decoding approach, we demonstrate that such suppression waves explain away ambiguous representation of stimulus position along the apparent motion trajectory. These waves thus preformat V1 population response for an unambiguous representation of the lrAM. Using an opponent motion energy approach, we demonstrate that the observed spatio-temporal pattern optimally encodes the stimulus velocity.

## Results

### Characterizing the mesoscopic spatio-temporal impulse response function

Two-step apparent motion sequences of various spatio-temporal characteristics (Fig 1, A and B) were presented to two behaving monkeys involved in a fixation task. The primary visual cortical response was measured at the level of the population using voltage-sensitive dye imaging (Amiram Grinvald and Hildesheim 2004; Chemla and Chavane 2010a). In response to a local stimulus (0.25° in diameter) presented for 100 ms in two different visual positions (separated vertically by 1° or 2°), activity arises at the retinotopic representation of these two positions and then spreads laterally over millimeters of cortical surface (Fig.1C: lower position, Fig. 1D: upper position) (A. Grinvald et al. 1994; Reynaud, Masson, and Chavane 2012; Muller et al. 2014). V1 activity is hereby reaching positions in space and time well beyond 1° and 50ms. As a consequence, the evoked spread covers a large cortical extent that can reach the representation of the other stimulus in space and beyond the inter-stimulus interval in time. The space- and time- constants of our responses were systematically quantified on the two monkeys and for the three stimulus durations we used (10, 50 and 100ms) on a 2D spatio-temporal (ST) map (Fig. 2A). To produce these ST maps, cortical activity was averaged within the apparent-motion trajectory (*dotted rectangle at frame 216 ms* in Fig. 1, C-G) to provide a unique spatial cortical dimension (ordinate in Fig. 2A). First, we extracted the space-constant of a gaussian spatial fit for all time points (see Fig. 2A, right-side of the maps). In both monkeys and across 19 sessions overall, the space-constant increased from 1.6 +/− 0.5 mm at response onset to reach a maximum of 3.3 +/− 0.2 mm, independent of the stimulus duration and monkeys (Fig. 2B, no significant difference observed between all stimuli durations, t-test with p>0.01). The time-constants of the response time-course at the central representation of the stimulus were measured using two halve gaussian functions fits (see Fig. 2A, below the maps). In both monkeys, the time-constant at response onset was on average 23.6+/− 17.2 ms for all stimuli durations (except for monkey BR with a mean value of 44.5 +/− 14.5 ms for 100 ms stimuli, see blue histogram in Fig. 2E), and 80 +/− 43.6 ms for response offset (Fig. 2F, no significant difference observed between all stimuli durations, t-test with p>0.01). Lastly, we also extracted the speed at which the response spreads across the cortical surface (see Fig. 2A, slanting lines) and obtained a distribution with peak values of about 0.26 +/− 0.14 m/s, similar across monkeys and stimulus durations (t-test with p>0.01), and similar to what has been observed in different species and states (Slovin et al. 2002; Sato, Nauhaus, and Carandini 2012; Bringuier et al. 1999; Reynaud, Masson, and Chavane 2012; Muller et al. 2014). This analysis showed that the spatio-temporal integrative properties of the primary visual cortex are mostly independent of stimulus duration and are able to cover a large spatial (3mm) and temporal (100ms) extent, bridging the cortical representation between our individual stimuli in space and time.

**Figure 1:**
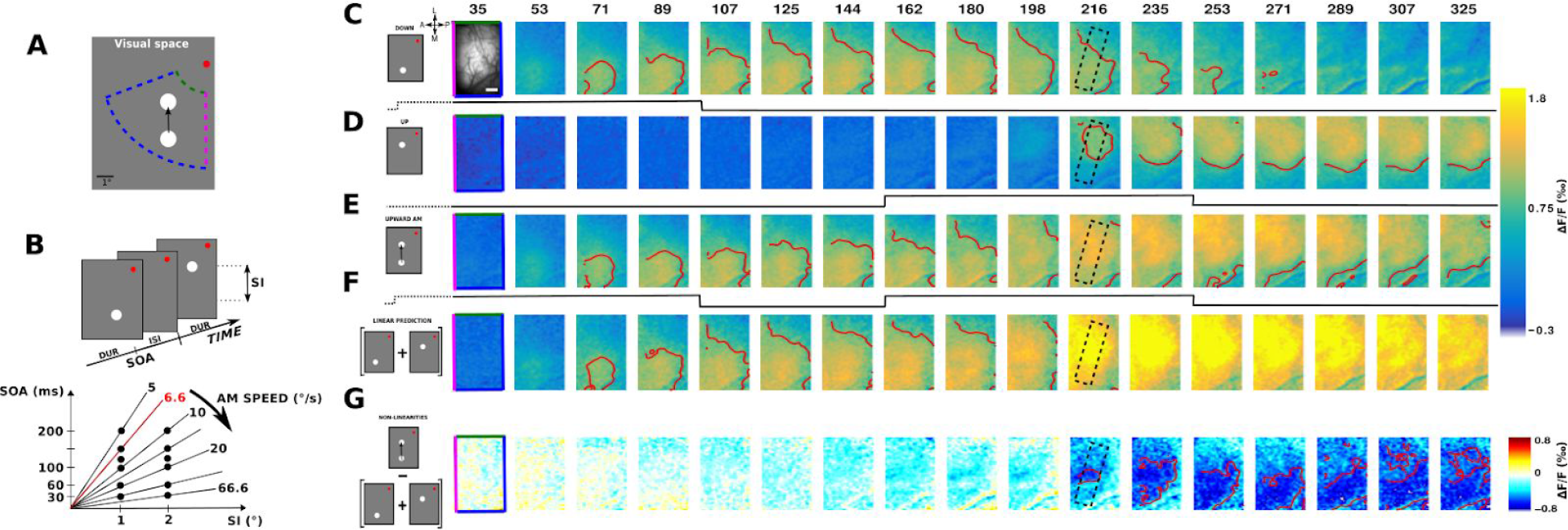
Experimental protocol and time-sequence of the cortical response to the long-range apparent motion (lrAM) **A:** Two-step lrAM stimuli are presented to two awake fixating monkeys in their bottom left visual field, while recording in their right visual cortex using VSDI. **B:** Spatio-temporal characteristics of lrAM stimuli, i.e. duration (DUR), interstimulus interval (ISI) and spatial interval (SI), were varied to cover a [5-66.6]°/s range of speed. **C-E:** Cortical representation of evoked VSDI activity as a function of time, in response to respectively, a 100 ms local stimulus in the down position, another one in the up position, and the sequence of these two stimuli (ISI = 50 ms and SI = 1°). The cortical area imaged is shown at upper left. The edge of the image color codes the retinotopic borders as represented in A such as the vertical meridian (magenta), eccentricities (green and blue). Scale bar: 2 mm; A: anterior, P: posterior, M: medial, L: lateral. Time in milliseconds after stimulus onset is shown at the top, while stimulation time is drawn at the bottom of each row (black lines). **F:** Activity pattern predicted by the linear combination in space and time of the response to stimulus 1 (row C) and the response to stimulus 2 (row D). **G:** Suppression pattern obtained by subtracting the observed apparent motion response (row E) and the linear prediction (row F). Red contours delimit amplitude activity above a certain threshold: 1 ‰ in panels C-F and −0.5‰ in panel G.

**Figure 2:**
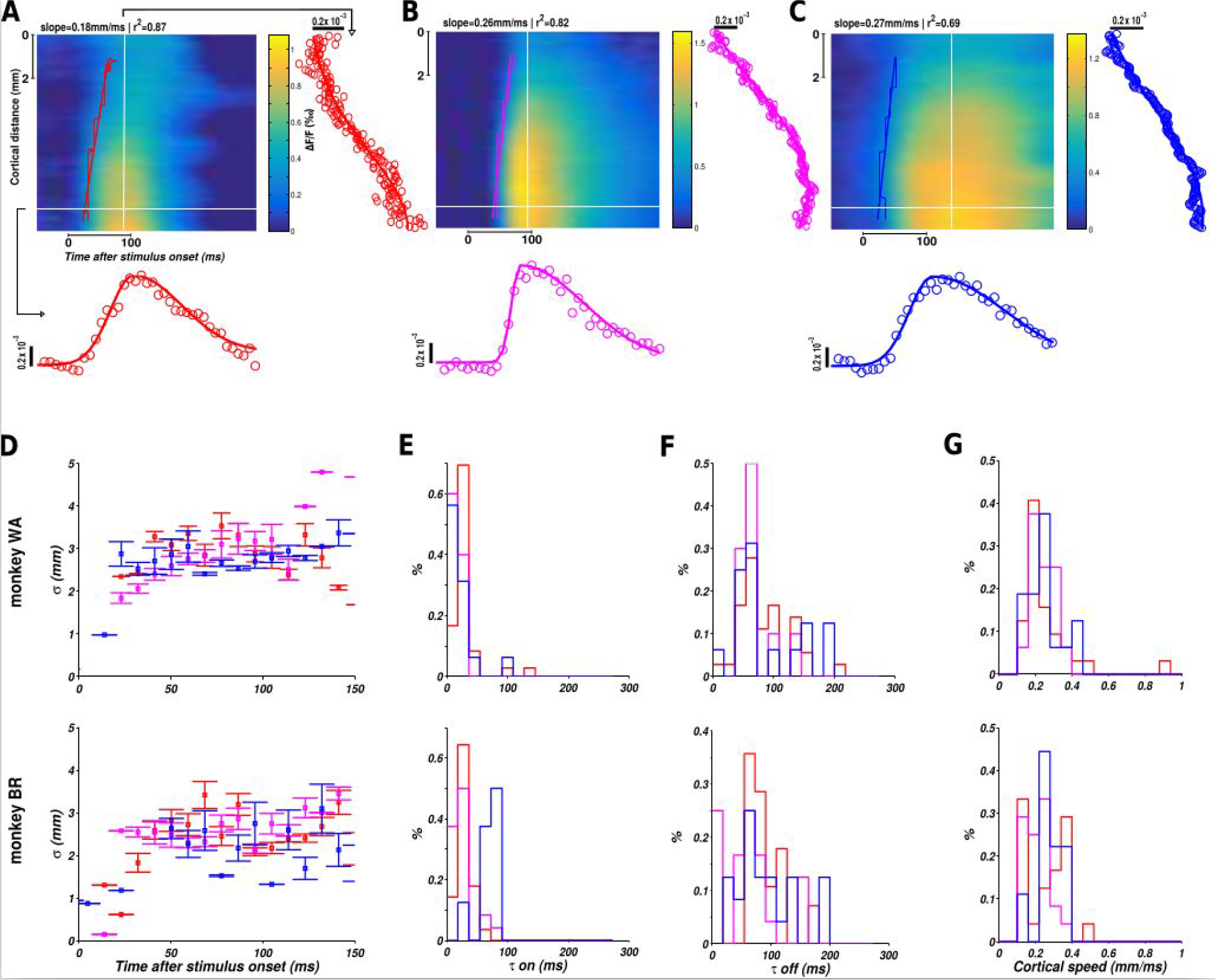
Spatio-temporal characteristics of cortical responses to a local stimuli. **A-C:**Spatio-temporal representations (ST) of the evoked cortical response to, respectively, 10 ms (A, red), 50 ms (B, purple) and 100 ms (C, blue) local stimuli. To produce the ST representation, we averaged spatial data along the stimulus trajectory (rectangle in frame 216ms, Fig1C-G). For each spatial point, the temporal data were fitted to a combination of two half Gaussians, as illustrated for one specific point in space (horizontal white line on the ST diagram) below the ST maps. Similarly, for each time frame, the spatial data were fitted to a Gaussian function as shown on the right side of each ST map for one specific point in time (vertical white line). **D:** Space-constant of the Gaussian spatial fit (sigma parameter) plotted as a function of time for the three considered durations (10 ms in red, 50 ms in magenta and 100 ms in blue) and for the two monkeys (top: monkey WA, bottom: monkey BR). **E:** Histograms of time-constant at response onset (**τ**on) estimated from the temporal fit of the response for the three considered durations and the two monkeys. **F:** Histograms of time-constant at response offset (**τ**off) estimated from the temporal fit of the response for the three considered durations and the two monkeys. **G:** Histograms of cortical speed of propagation estimated by linear regression on response latency (stairs-step contours, slanting lines and slope of the linear regression) for the three considered durations and the two monkeys.

### The evoked response to the lrAM is shaped by a suppressive wave

We next asked whether such lateral interactions contribute to shape the evoked population response to the temporal succession of these two stimuli. For that purpose we measured the cortical population response to a two-stroke upward apparent motion sequence (Fig. 1E). Such temporal sequence generates a propagation of activity starting at the cortical representation of the first stimulus (S1) and moving to the cortical representation of the second stimulus (S2), a cortical correlate of the illusory motion (Jancke et al. 2004). The observed pattern of activity departs from the pattern predicted by a simple linear summation of the lower and upper stimuli (Fig. 1F). If we subtract the observed (Fig. 1E) and the linear predicted responses (Fig. 1F), two deviations from non-linearities are observed. First, a suppression emerges at response onset and at the cortical representation of S2 (compare 1D and 1G at frame 216ms). The suppression then gradually propagates over the cortical surface towards the representation of S1 (Fig. 1G). We can hypothesize that the evoked activities by the two stimuli composing the lrAM sequence interact together to generate this dynamic pattern of suppression. Since the suppression is observed at the onset time of the response to S2, it has to be due to the activity dynamics generated by S1 interacting with the integration of S2. However, the propagation of suppression from the representation of S2 towards the representation of S1 is probably due to the activity dynamics evoked by S2 interacting with the residual activity evoked by S1. Therefore, the suppression wave could likely be the result of multiple interactions (e.g bidirectional) between the activities evoked by the stimulus sequence.

### The suppressive wave is systematically observed

To better investigate how spreads of evoked activity and suppression shape the representation of lrAM, we first show ST representations of examples taken for both monkeys and three stimuli speeds. The example of Figure 1 is shown in Figure 3A (6.6°/s). In these ST representations, we can observe a clear propagation of activity in response to a local stimulus (slanting lines in Fig. 3, A and B) that is remarkably similar across both monkeys (Fig. 3, A and B, *first rows*) and speeds (three columns respectively for 6.6°/s, 10°/s and 33.3°/s, as shown in Fig. 2F). The ST representation of non-linearities (lower rows) recentered on S2 onset, shows that suppression first appears at the cortical representation of S2 and at S2 response onset, and then propagates towards the representation of S1, at a similar speed than the one observed for the evoked activity to the first stimulus (Fig. 3, A and B, *second rows, slanting lines*). In both monkeys and the three examples shown, this suppression propagates in a direction opposite to the apparent motion sequence, from S2 to S1 representations. Functionally it results in silencing the residual activity generated by S1.

**Figure 3:**
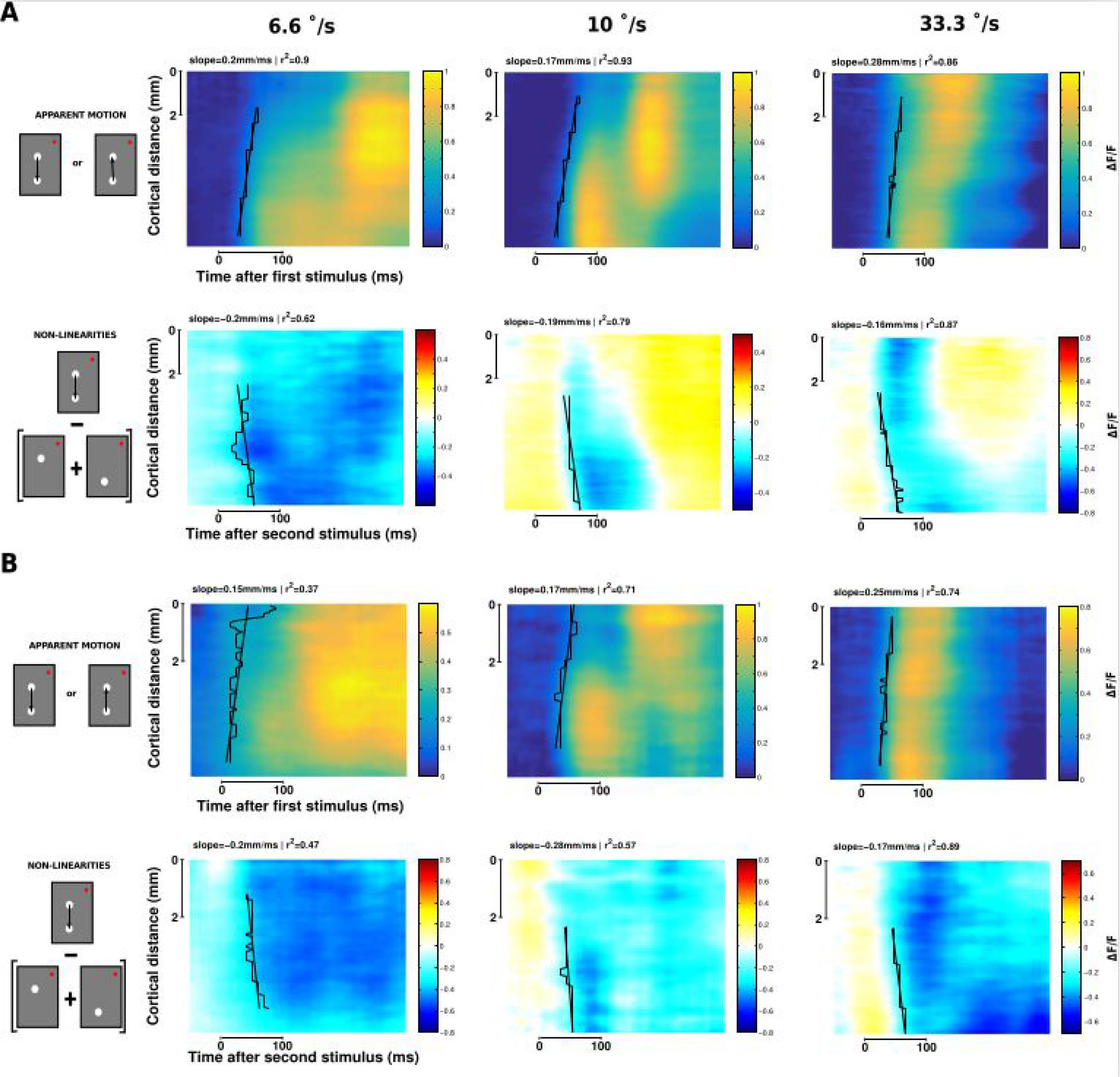
The apparent motion stimulus induces a systematic suppression wave. Spatio-temporal representation of VSDI responses to two-stroke apparent motion stimuli for three different speed (6.6°/s, 10°/s and 33.3 °/s) and two animals (**A:** monkey WA, **B:** monkey BR). The upper rows of A and B represent the observed response and the lower rows the non-linearities of the response (observed - linear prediction). Estimates of speed propagation are reported on each ST diagram (black stairs-step are contours at threshold level, slanting lines are the slope of the linear regression). Similar values are observed for both the observed activity and the non-linearities.

### The suppressive wave propagates at the same speed and with same extent as the evoked spread

This suppressive wave was systematically observed for all two-stroke lrAM conditions tested (see Fig.1B). This can be seen in the ST evoked response (centered on the onset of S1) and nonlinearities (centered on the onset of S2) averaged across all conditions and sessions for both monkeys (Fig. 4A). To better understand the origin of the suppression dynamics, and its dependence on stimulus conditions, we characterized its spatio-temporal properties. First, we measured the onset of the apparition of the suppression at S2 position. The latency of the observed suppression was the same as the latency of the activity evoked by S2 alone (Fig. 4B, respectively 39.5 +/− 2.0 ms vs. 38.6 +/− 1.6 ms for monkey WA and 36.6 +/− 1.8 ms vs. 36.9 +/−2.1 ms for monkey BR, non-significantly different, t-test with p = 0.77 and p = 0.35 respectively for WA and BR). However, the suppression resulted in significantly delaying the response onset evoked by S2 when presented within the apparent motion sequence (54.2 +/− 2.0 ms and 68.3 +/− 5.3 ms for WA and BR respectively, Fig. 4B). Then, we quantified the spatial extent of the suppression (**σ** of a Gaussian fit, Fig. 4C). In all conditions, the spatial extent of the suppression was of about 2.8 mm (2.49 +/− 0.14 mm for WA and 3.08 +/− 0.18 mm for BR), similar and non significantly different than the spatial extent of the evoked response (2.99 +/− 0.11 mm and 2.41 +/− 0.17 mm for WA and BR respectively). Thus the suppressive wave starts at similar latency and covers similar spatial extent. We next characterized the speed of propagation of activity (Fig. 4D black) and suppression (Fig. 4D blue), plotted as a function of stimulus speed. Remarkably, on both monkeys, the observed cortical speeds were identical for both the propagation of activity and the suppression and completely independent of the lrAM speed (0.28 +/− 0.26 m/s and 0.27 +/− 0.4 m/s respectively for WA and 0.21 +/− 0.15 m/s and 0.27 +/− 0.2 m/s respectively for BR). However, from the ST plots in Figure 3, we noticed that the suppression does not seem to spread but rather propagates as a wave (Muller et al. 2014, 2018). To probe for this hypothesis we thus compared the dynamics of the response peak position (**μ** of a Gaussian fit). In a spread, typically, the response peak will not move in space, as observed for evoked response (Fig. 4E, the peak spatial position is not changing with time, slope of −1.3×10^−5^ +/− 1.1×10^−4^ m/s and 1.6×10^−4^ +/− 3.4×10^−4^ m/s for WA and BR respectively), whereas in a wave it will follow the onset spatial displacement, which is what we found for the suppression (Fig. 4E, the peak moves from position 2 to position 1, negative slope of −0.05 +/− 0.007 m/s and −0.034 +/− 0.005 m/s for WA and BR respectively). Altogether, our results show that the suppression is initiated at response onset, have similar spatial extent and propagation speed as the activity evoked response. Furthermore, although evoked activity are waves hidden by spatial averaging (Muller et al. 2014), the suppression is still seen as a wave in the averaged data. This strongly suggests that the suppression is likely to be mediated by the same general process generating the propagation of evoked activity, most probably the intra-cortical horizontal network (Muller et al. 2014). If the suppression is generated along the propagation of activity, one prediction is that it should decrease in strength with spatial and temporal separation between the two stimuli composing the lrAM. This is indeed what was observed, the suppression strength decreases as a function of stimulus onset asynchrony and spatial separation (Fig. 4F, t-statistics on the slope of the linear regression gives t = −0.92 with p=0.18 and t = −6.3 with p = 3.6×10−6, respectively for a spatial interval of 1⁰ and 2⁰ (WA); t = −1.2 with p=0.12 and t = −1.6 with p = 0.05 (BR)).

**Figure 4:**
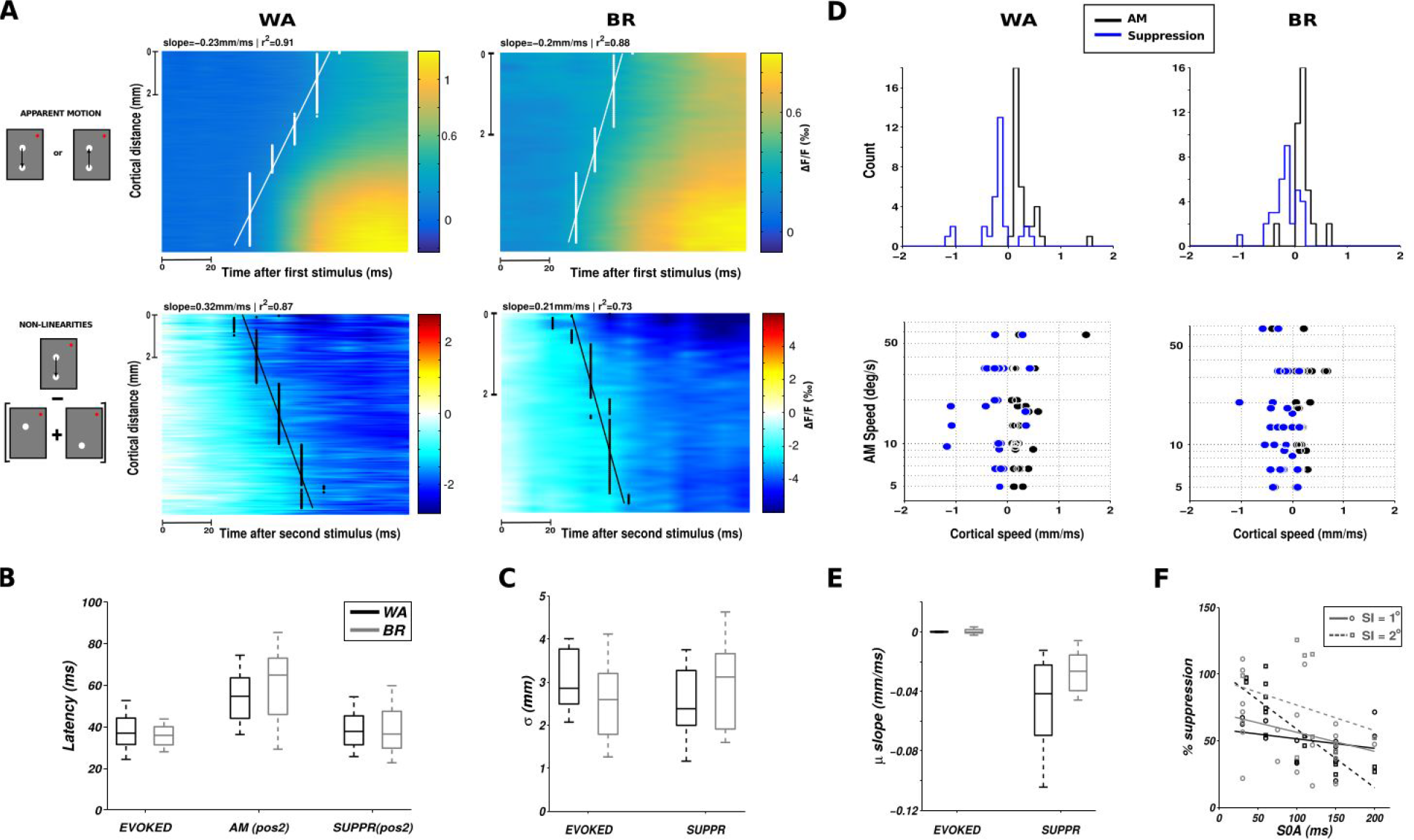
The suppressive wave has the same properties as the evoked intra-cortical propagation. **A:** Spatio-temporal VSDI activity (top row) and non-linearities (bottom row) averaged across all lrAM speed conditions and centered on stimulus 1 (S1, top row) or stimulus 2 (S2, bottom row) onset, for both monkeys (columns). **B:** Boxplot of latency estimates comparing the onset of activity evoked by S2 alone (“evoked” condition), the response onset evoked by S2 when embedded in the lrAM sequence (“lrAM” condition) and the onset of the suppression at S2 position (“suppr” condition). Boxplots illustrate median, 25 and 75% quartiles and minimum and maximum of the distributions across all lrAM speed conditions, for the two monkeys (black WA, gray BR). **C:** Boxplot of space-constants (parameter **σ** of a Gaussian spatial fit) comparing the evoked response and the suppression, for the two monkeys. **D:** For each condition in both monkeys (columns), we estimated the speed of propagation of the VSDI (black) and the non-linearity (blue). The upper row shows frequency histograms and the lower row these speeds as a function of the speed of the lrAM stimulus. **E:** Boxplot of the response peak propagation speed (slope of the linear regression on the parameter **μ** of a Gaussian spatial fit) comparing the evoked response and the suppression, for both monkeys. **F:** Suppression strength (normalized to the maximal response activity) as a function of stimulus onset asynchrony and spatial interval (open circle for SI = 1⁰, open square for SI = 2⁰), for both monkeys (black WA, gray for BR).

### The suppressive wave can be the result of a dynamic gain control

What can be the origin of such suppressive wave? Since inhibitory intra-cortical axons have more limited spatial extent (Buzás et al. 2001), and that feedback from higher areas are excitatory (Salin and Bullier 1995), we can hypothesize that it does not result from a simple net inhibition, but rather as a byproduct of the excitatory/inhibitory balance (Tsodyks et al. 1997; Ozeki et al. 2009). Indeed, as demonstrated using center-surround stimulations, the suppressive wave can be the result of a simple dynamic input normalization fed by propagation along the horizontal network (Reynaud, Masson, and Chavane 2012). To determine the possible mechanisms generating the observed suppression, we used a mean-field model designed to reproduce accurately VSDI (Zerlaut et al. 2018). In this model, it was assumed that each pixel of the VSDI represents the average Vm of two populations of interacting neurons, excitatory regular-spiking (RS) neurons, and inhibitory fast-spiking (FS) neurons (Chemla and Chavane 2010b). Arranging this model into a spatially extended interconnected populations of RS-FS cells (Fig. 5A, see Methods) allows to simulate the propagating waves observed in awake monkey under VSDI. The great advantage of such model is to explicitly take into account conductance-based interactions (COBA) as well as a different gain between excitation and inhibition. These ingredients are often neglected as they introduce difficulties in mathematical tractability of mean field models (Landau et al. 2016; Vogels, Rajan, and Abbott 2005). Nevertheless, these features are biologically relevant and, as we show here, are actually the main elements determining waves suppression. Examples of two independent waves are shown in Fig. 5B (upper row). When the two stimuli are presented in succession (see Fig. 5B lower left) the observed response shows a suppression (Fig. 5B lower right), whose values are quantitatively similar to those of experimental data (suppression of around 50% of the response max). Such suppressive effect was robustly observed across a wide range of the parameters space. The first parameter that was found to strongly affect the suppression is the ongoing spontaneous activity of the system pre-stimulus. As we report in Fig. 5C (COBA model, red dots), the suppression decreases when the spontaneous activity of the system increases (see example marked by a circle, Fig 5D). Moreover, two further mechanisms were necessary to explain this suppressive effect. First, inhibitory cells need to have a higher gain than excitatory cells. When the gain of FS cells was reduced (see inset of Fig. 5C) to have a gain closer to the one of RS cells, the suppression effect was strongly affected (blue dots in Fig. 5C, example marked by a square in Fig. 5D). Accordingly, increasing FS cell gain (cyan dots in Fig. 5C, example marked by a pentagon in Fig. 5D) increases the suppression strength. Second, the interaction between excitatory and inhibitory inputs needed to occur through conductances-based mechanisms. Indeed, when using a current-based (CUBA) model (see Methods), we mostly observed facilitation (black triangles in Fig. 5C) that do not appear to propagate (see example marked by a triangle, Fig. 5D). While we do not exclude that such suppression may be observed in current-based synapses, it is clear from these data that the non-linearity of voltage dependent synapses induces a strong suppression in VSDI signal. The suppression can thus be explained by the mesoscopic combination of the nonlinearity of conductance interactions and the differential gain of excitatory and inhibitory cells.

**Figure 5:**
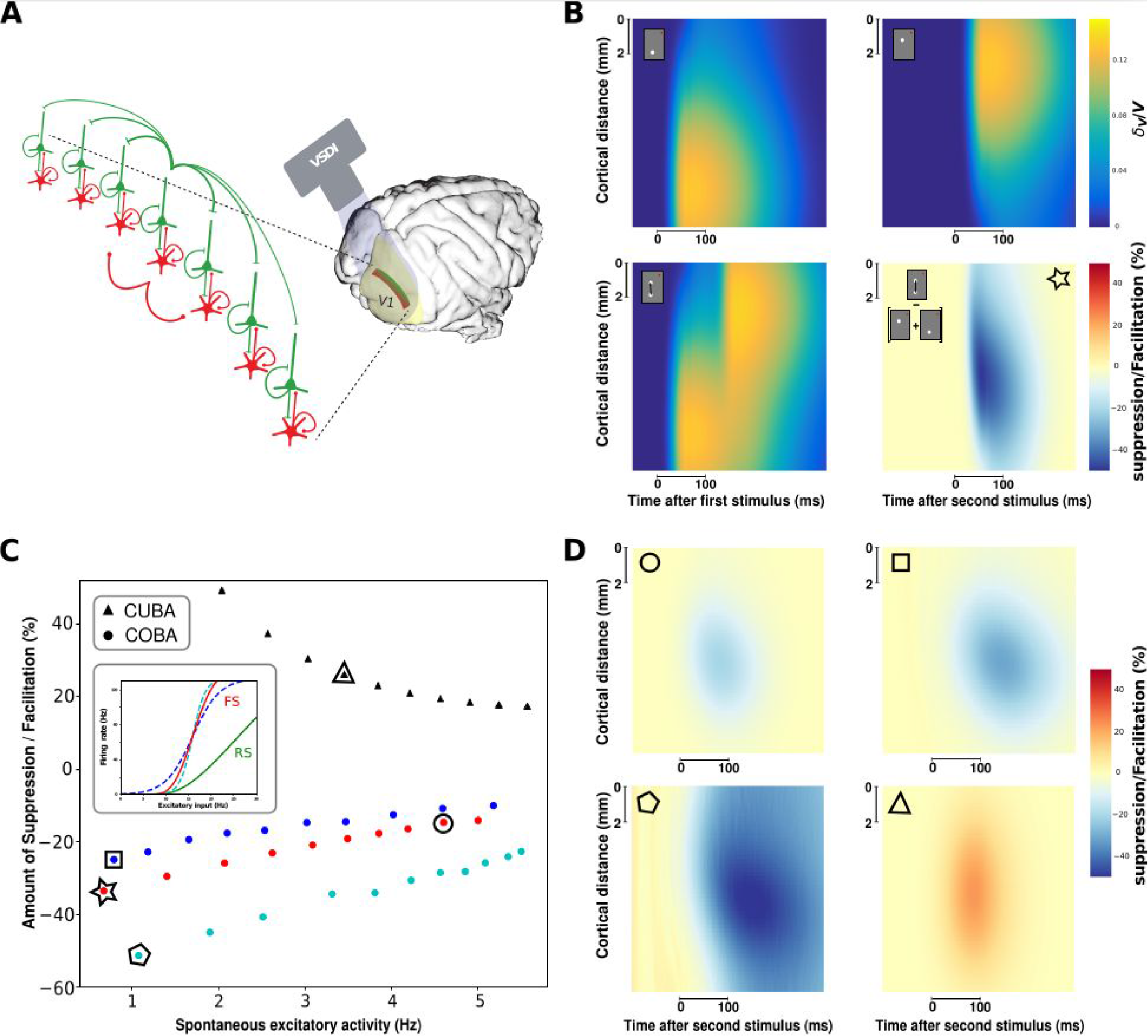
A computational model to investigate the possible origin of the suppressive wave. **A:** Mean-field model of excitatory and inhibitory neurons distributed on the cortical trajectory of the stimulus with horizontal connectivity (longer for excitatory than inhibitory neurons). **B:** Model ST response to the first stimulus (upper left), the second (upper right), the apparent motion sequence (lower left) and the non-linearities normalized to the maximal response over space and time of the response to single stimuli (lower right). The input has an amplitude ν0=20 Hz. **C:** Amount of suppression/facilitation as a function of the spontaneous excitatory firing rate. Colored dot stand for different interneurons gain (see inset), while black triangles stand for the Current-based (CUBA) model, that shows little suppression but facilitation. The input has an amplitude ν0=10 Hz. **D:** Representative ST suppressive/facilitative patterns as marked in C by different geometric shapes (circle, square, pentagon, triangle). The star in C corresponds to the model parameters used for obtaining the suppressive pattern shown in B.

### The function of the suppressive wave is to explain away ambiguous representations

What can be the function of the suppressive wave? Here we propose that it will shape an unambiguous representation of motion along the apparent-motion trajectory. Indeed, silencing the cortical representation of the initial stimulus when the second stimulus is being processed will have as a consequence to represent only one stimulus at a time, hereby improving motion representation by explaining away ambiguous position representation (problem of “phenomenal identity”) (Ternus 1926). To quantify such hypothesis, we developed a simple algorithm to decode, at every instant, what is the most probable stimulus position that evoked the observed cortical spatial profile out of four categories: no stimulus, S1, S2, or joint S1 & S2. We used the ST representations of the evoked activity to the apparent motion sequence (Fig. 6A) and used the linear prediction (Fig. 6B) as a control. The decoding was computed using the joint probability that the spatial profile observed at one point in time (white profile) is drawn from the spatial profile observed during blank (first row, black), S1 (second row, red), S2 (third row, blue), or the joint S1 & S2 (last row, green). In the example shown in figure 6, we apply this decoding method to the activity evoked by a 6.6°/s two stroke apparent motion stimulus (Fig.6A). When S1 is presented (red), the probability that the spatial profile of the evoked response will be similar to the blank distribution is quickly dropping from 1 to 0 and the probability that the evoked response will be decoded as being evoked by S1 alone is jumping from 0 to 1 very rapidly (in 10ms). When S2 is presented (at time 50ms) there is a sharp and rapid transition from the evoked activity being decoded as S1 to S2 (blue) in about 50ms. However, the probability that the evoked activity is evoked by S1 & S2 at the same time (green) is only increased moderately (peaking at 0.5) and transiently. In contrast, when we apply the same approach to the linear prediction (Fig.6B), while the beginning of the decoding is the same (two first rows), as expected, when S2 appears, the evoked activity is ambiguously decoded as being attributed to S2 or S1 & S2 conjointly with similar probability (around 0.5).

**Figure 6:**
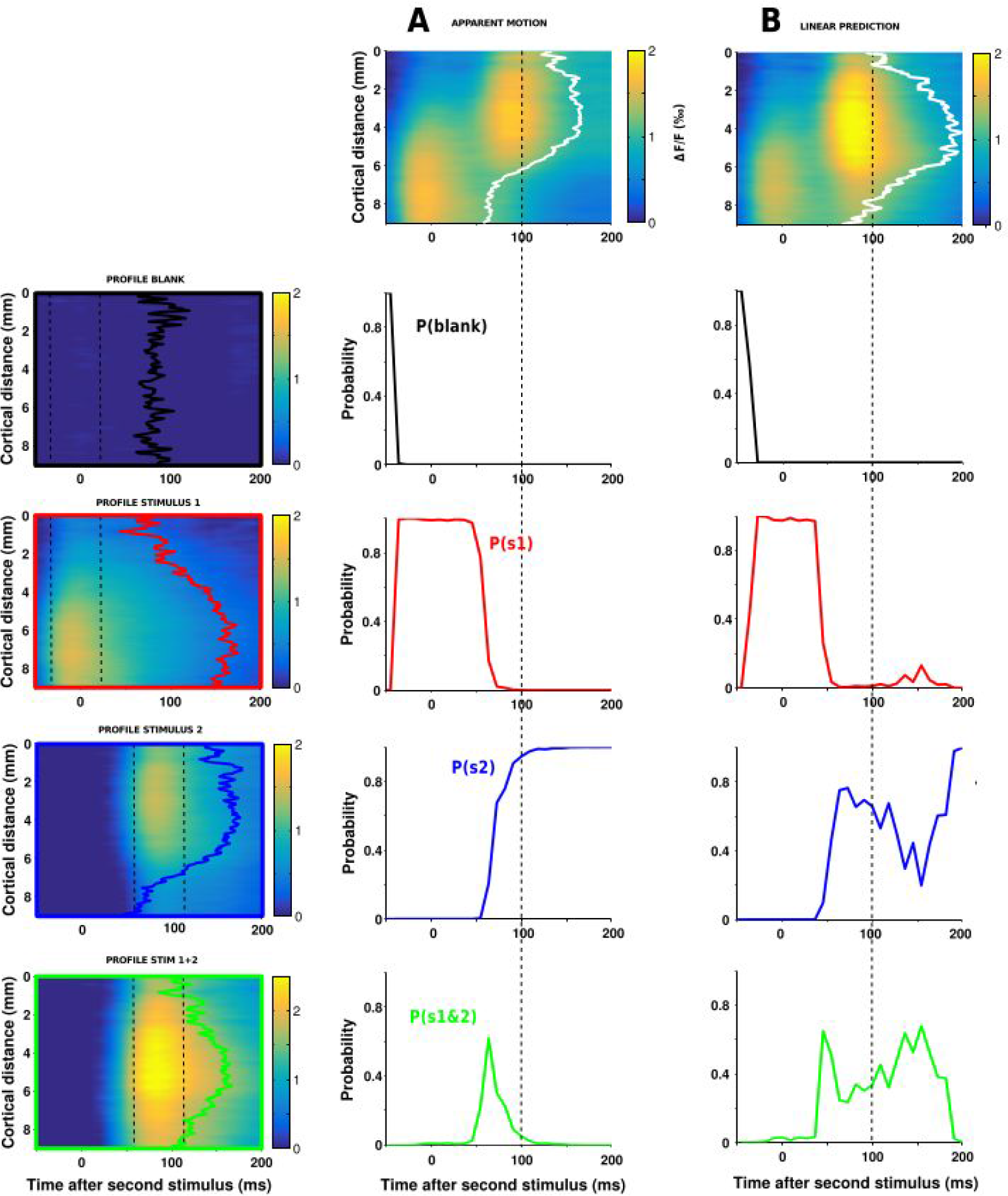
A dynamic decoding of stimulus position: Principle. The decoding of stimulus position on ST maps, here taking the example of the activity evoked by a 6.6 °/s lrAM stimulus shown in **A** or the activity pattern predicted by the linear combination in space and time of the responses to both individual stimuli in **B.** The decoding consists in evaluating the probabilities that the spatial profile observed at each point in time (white contours in A and B) is similar to one of the four spatial profiles shown on the left column: Blank (first row, black profile), S1 (second row, red profile), S2 (third row, blue profile), and the joint S1 & S2 (last row, green profile). Each profile was computed by averaging the corresponding ST response in a 50ms-window around the time of maximum response and normalized. The four color-coded probabilities are respectively plotted as a function of time (time 0 corresponds to the onset of S2) for the lrAM response (**column A**) and for the linear prediction (**column B**). Compared to the linear prediction, the actual signal is more rapidly decoded, revealing a likely function of the suppressive wave: shaping stimulus position representation.

We applied this approach to all speeds and sessions in both monkeys (Figure 7A&B), for spatial interval of 1°, differentiated across the different inter-stimulus intervals (ISI). We separated these conditions because, when S2 appears, the residual activity in response to S1 will be less important for long ISI (the offset time constant being of the order of 80 ms). In both monkeys and for ISI <= 50ms, the averaged results confirm the individual example shown in Figure 6: the evoked activity results in a sharp and clear transition from the representation of S1 to the representation of S2, with only transient increase of the representation of S1 & S2 conjointly. In comparison, the linear prediction always leads to an ambiguous representation that cannot tease apart the probability that the evoked activity is coming from S2 alone or S1 & S2 together (blue and green curves merging together). For an ISI >= 100ms, in contrast, the evoked activity resembles more the linear prediction, as expected.

**Figure 7:**
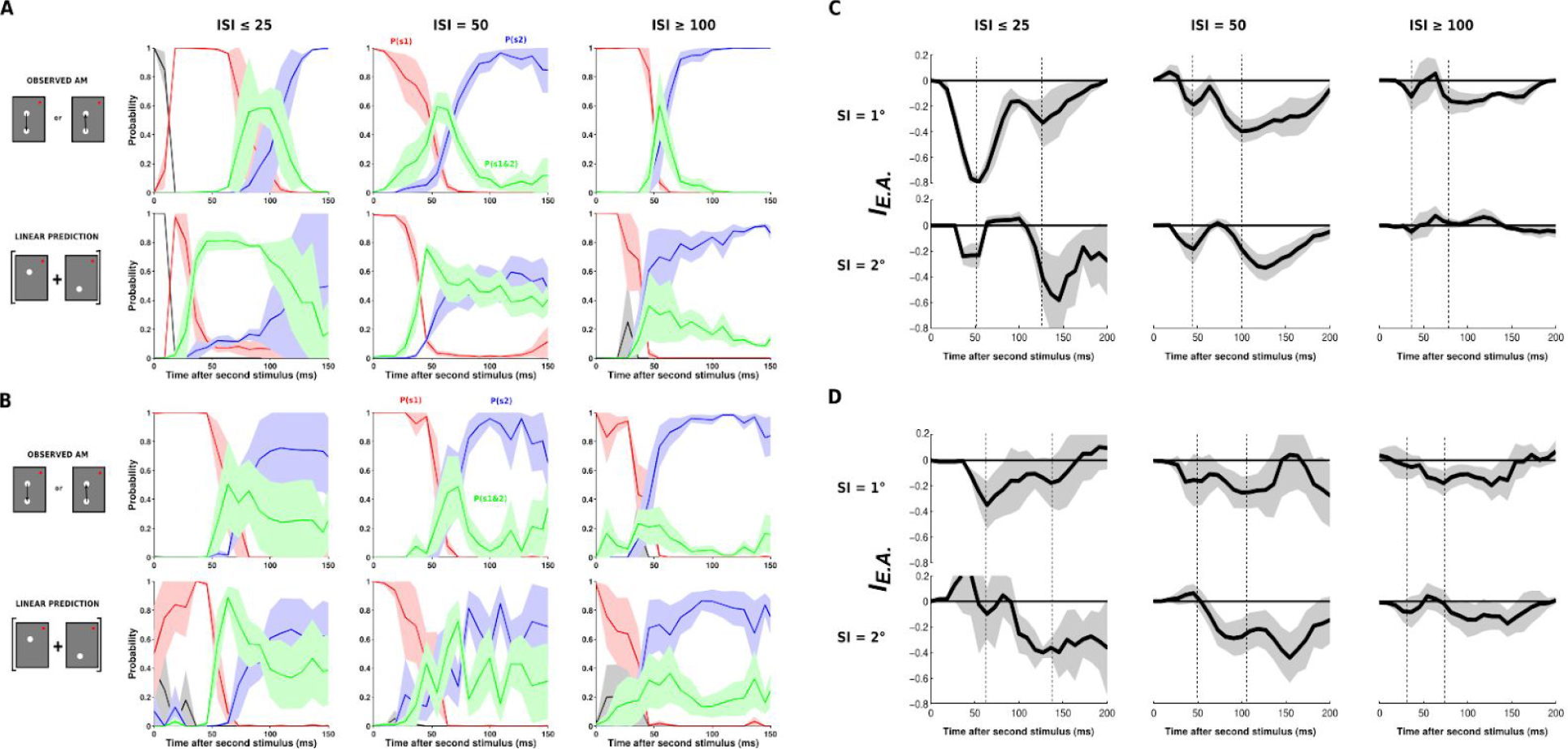
A dynamic decoding of stimulus position: Application to all lrAM speeds and sessions. **A:** Color-coded probabilities (same as Figure 6) for the observed lrAM response (first row) and its corresponding linear prediction (second row) for monkey WA, averaged across three ISI categories: ISI < 25 ms (left column), ISI = 50 (central column) and ISI > 100 ms (right column). **B:** Application of the decoding algorithm to all the data of monkey BR. **C:** Explaining away index (see text and methods) computed as the probability of detecting joint S1 & S2 in the observed response minus the probability of detecting joint S1 & S2 in the linear prediction, from monkey WA data shown in panel A. **D:** Explaining away index from monkey BR data shown in panel B.

To quantify the effect of explaining away ambiguous positional representations during lrAM stimulations, we calculated an index by subtracting the probability of detecting joint S1&S2 in the observed and the linear prediction for both monkeys, 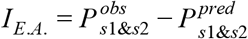 (Fig. 7C&D), and both stimuli spatial intervals (SI) of 1 and 2° (first and second rows respectively). In all conditions but the long SI and long ISI, a systematic decrease of the index was observed. This reveals a dynamic effect of explaining away the ambiguous representation of S1&S2. Importantly, in both monkeys and practically all conditions (ISIs and stimulus separation), we observed two peaks in the index decrease. They correspond to the bidirectional interactions occurring for each of the two evoked waves. The first peak corresponds to the effect of delaying response onset to S2 (by propagating activity from S1 to S2), and the second peak corresponds to a shortening of the representation of S1 (by propagating activity from S2 to S1). Importantly, this calculation revealed two further phenomena that are expected because of the propagation delay and spatial extent. First, the timing of the second peak is delayed when going from 1 to 2° spatial separation. Second, the general amplitude of the decrease diminishes from short to longer ISI.

### Unambiguous representation for optimal encoding of velocity in V1

Shaping the cortical population representation of the lrAM could promote an accurate encoding of direction-selective motion signals for an optimal read-out by downstream area. To test whether the measured cortical response encodes an accurate direction-selective signal, we applied opponent motion energy filters directly to V1 population responses (Adelson and Bergen 1985). Indeed, direction selectivity in MT is well described and captured by motion energy models (Adelson and Bergen 1985; Rust et al. 2006). Such an approach is generally developed to model MT receptive field from a spatio-temporal input image. The rationale here is to apply the same processing directly to V1 population responses that feed downstream areas such as MT or V4. This is justified by the fact that the cortical extent imaged here (~ 9mm, corresponding to 3°, see Dow et al. 1981; Van Essen, Newsome, and Maunsell 1984) actually corresponds to the V1 cortical extent converging to a MT or V4 neuron at our recorded eccentricity (3°, see Albright and Desimone 1987; Gattass, Sousa, and Gross 1988). Since we record VSD responses that represent both sub- and supra-thresholds activities (Chemla and Chavane 2010b), we first processed our ST maps through a non-linearity to account for the VSD to spike rate transformation (Chen, Palmer, and Seidemann 2012) (Fig 8A). The resulting ST maps were convolved with a set of spatio-temporal filters covering a wide range of speeds and scales. For a given value of filter speed and scale, we squared and summed the convolution from filters in quadrature, and subtracted the resulting phase-independent measure of local motion energy for opposite directions (ie. MEu - MEd) to obtain the opponent motion energy response (OME, Fig 8A). We thereby obtained the opponent motion energy for all speeds, scales and directions. For each position on the ST map, we could hence extract the filter velocity for which the opponent motion energy is maximal, that we represented for both monkeys, and different velocities (10°/s upward in monkey1, Fig8B and −33°/s downward in monkey 2, Fig8C). In this representation, the color hue represents the velocity of the filter yielding a maximal opponent motion energy and the color intensity its amplitude (as a fraction of the maximum evoked fluorescence response). The contour of the evoked response is overlaid in white to ease comparison. The same analysis on the corresponding linear predictions serves as a control. For all the conditions we explored, we then extracted the values of the optimal velocity within a ST region of interest (between S1 and S2’s centers and from 10 to 200 ms after stimulus 2 onset) and represented them as a function of the AM speed for both monkeys (Fig 8D and 8E). Our results show that the ST response, shaped through the suppressive wave, is indeed generating a direction selective motion energy for a speed that is well correlated with the stimulus speed. In other words, intra-cortical non-linear interactions in V1 promote an unambiguous optimal encoding of velocity-selective motion signal along the apparent motion path.

**Figure 8:**
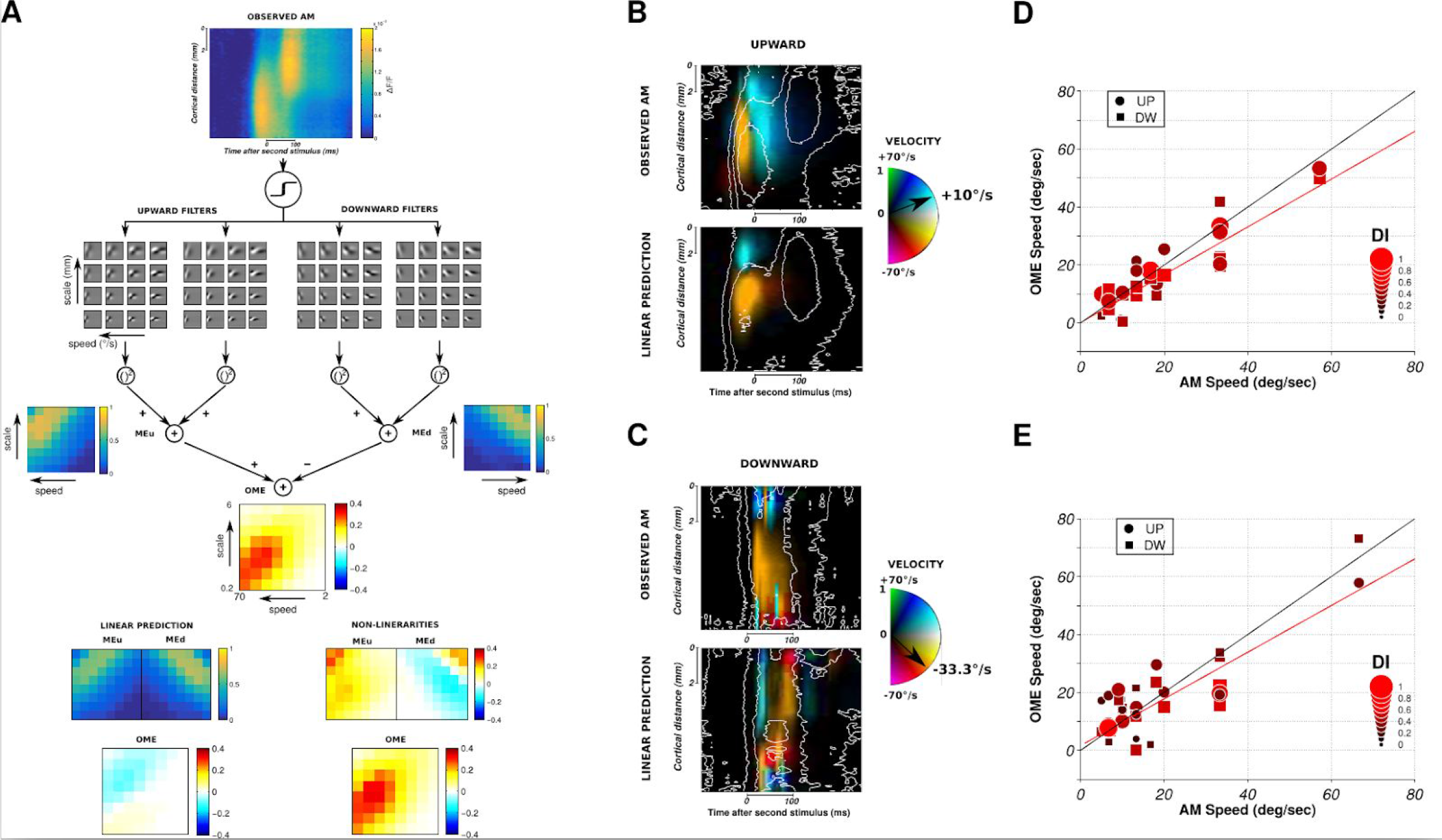
Encoding of direction-selective motion signal. **A:** Application of the opponent motion energy model (Adelson & Bergen, 1985) to the ST representation of cortical response to an upward 10° /s AM sequence shown at the top. The first step consists in convolving the ST data with a set of oriented ST filters. Phase-independency is obtained by squaring and summing the outputs of quadrature pair of filters, while motion opponency is obtained by subtracted the two oriented motion energies (OME = MEu-MEd). The maximal energy values for each ST filter are plotted as a function of speed (in °/s) and scale (in mm). The energy values resulting from the same computation applied on the linear prediction and the non-linearities for this AM sequence are respectively shown at the bottom left and right. **B:** ST representation of the opponent motion energies computed in panel A. For each ST position, the filter velocity for which the energy was maximal is represented as different color hue. The amplitude of the energy is coded as color intensity. For comparison, the result for the corresponding linear prediction is shown below. Non-linearities correspond to observed AM - linear prediction. **C:** Same than B for monkey BR, for another AM sequence condition (33.3° /s downward motion). **D:** Filter speed that generated the strongest OME within a ST region of interest (see Methods) as a function of the actual lrAM speed for monkey WA. The color and size of the dots (upward motion conditions) and squares (downward motion conditions) code for the value of the direction-selectivity index (DI). **E:** Same than D for monkey BR.

## Discussion

We have shown that intra-cortical interactions play a key role in shaping the sensory representation of the long-range apparent motion within the retinotopic map of V1 in awake monkeys. Our results demonstrate that intra-cortical propagation encompasses large spatial and temporal distances allowing to link information between stimuli presented in distal spatial positions (spatial constant of about 3 mm, equivalent to 1°, and time constant of about 80 ms). Importantly, above these values, the apparent motion illusion gradually fades out (Kolers 1972; Cavanagh and Mather 1989). In response to a two-stroke lrAM sequence, we observe a clear displacement of activity on the cortical surface that deviates from the linear prediction in two aspects. First, the initial stimulus suppresses and delays the response to the second stimulus. Second, a suppressive wave is evoked by the second stimulus that strongly and rapidly attenuates the residual activity evoked by the first stimulus. The spatio-temporal characteristics of the suppression show similar spatial constant and similar propagation speed as what was observed for the evoked activity, independent of the speed of the apparent motion stimulus. However, the suppression propagated as a true wave in direction of the initial stimulus position, even at the trial-averaged level, an observation that departs from what we observed in the evoked activity (Muller et al. 2014). We propose that the suppression arises from a simple gain-control mechanisms pooling feedforward and horizontal inputs (Reynaud, Masson, and Chavane 2012). To demonstrate this, we used a conductance based mean-field model developed to account for VSD dynamics (Zerlaut et al. 2018). This model shows that the observed suppression can be explained by nonlinear conductance interactions, combined with the different gain of excitatory and inhibitory cells. A decoding approach demonstrates that the suppressive wave acts as explaining away the ambiguous representation allowing to represent only one stimulus at a time in the cortex. Using opponent motion analysis applied to the population response, we demonstrate that such unambiguous representation allows V1 to encode accurately the velocity signal of the lrAM that could support the read-out process from downstream areas.

### Suppression and normalization as generic operations in the visual system

The dynamics of the suppression is seen here as a central and key mechanism by which the input is shaped and normalized by V1 populations. When more than one stimulus is present in a visual scene, suppressive interactions between the feedforward-driven activities is what is traditionally reported, such as the well documented surround suppression (Blakemore and Tobin 1972; Angelucci et al. 2002; Cavanaugh, Bair, and Movshon 2002). This suppression is generally attributed to be an emergent property of the divisive normalization computation (Carandini and Heeger 2011). Importantly, we have shown that this normalization process is dynamic and propagate from the representation of the stimulus surround towards the representation of the center (Reynaud, Masson, and Chavane 2012). Adding a new lateral input (mostly excitatory at long-distance) therefore results in a decrease from the linear prediction, a paradoxical inhibitory effect (Tsodyks et al. 1997) well captured by Stabilized Supralinear Networks (Ozeki et al. 2009). Similar suppression was also seen in response to the line-motion stimulus (Jancke et al. 2004), however, in that stimulus conditions, it was preceded by a transient facilitation. The main difference with our paradigm is that, in the line-motion condition, the second stimulus, a bar, provides a feedforward activation all along the trajectory of the evoked wave. In the apparent motion, the interactions involve only cortical interactions at positions that do not receive any feedforward input. This may explain the differences observed with the line-motion stimulus. We believe that dynamic non-linear interactions subtended by intra-cortical network acts as a general gain control shaping the representation of visual stimulus in space and in time.

### Modeling the suppressive waves

Possible mechanisms underlying the observed suppressive effects were investigated using a mean-field computational model (Markounikau et al. 2010), that has been spatially extended. We found that the model can reproduce the observed suppression, provided two mechanisms are present: excitatory and inhibitory cells have a different gain, with a higher gain for inhibition, and excitatory and inhibitory synaptic inputs must combine through conductance-based interactions. Although these two mechanisms are well known, they are usually neglected in mean-field models because they represent a mathematical difficulty. The classic mean-field models with linear (current-based) interactions and uniform gain in all cells, fail to reproduce the suppressive effect of propagating waves, and thus the present model can be considered as a step towards biologically more realistic mean-field models. Hence, by constructing a realistic mean-field model, we could demonstrate that this suppression wave is an expected byproduct of the known anatomy and does not need to be expressed solely by pure inhibition. This computational approach demonstrates how excitatory and inhibitory propagation of activity along horizontal network can dynamically change the cortical gain control resulting in the emergence of the observed suppression dynamics.

### Backward suppression to keep track of object identity along the apparent motion path

This suppression can help to represent unambiguously one object at a time on the cortical surface, as our decoding model suggests. This means that the lateral interactions can link the transient spatio-temporal events while keeping track of the object moving along the trajectory. This could be a first mechanism involved in solving the correspondence problem (Ullman 1978). This problem, first introduced by Ternus as a problem of phenomenal identity (Ullman 1978; Ternus 1926), put to light the fact we need to keep track of the identity of an object in movement, and, in the case of multiple objects present at each time frame, a problem of correspondence may occur. The literature clearly show that the correspondence is solved through spatio-temporal coherence more than shape or color consistency (Kahneman, Treisman, and Gibbs 1992). The correspondence, called “reviewing” by Kahneman et al. (1992) was proposed by these authors to “*operate(…) backward, (…) select(…) only a single item, and (…) is guided mainly by the features that control the unity and continuity of an object over time, but not by the shape, color, or content of the target*.” We believe that the mechanisms of backward suppression demonstrated here is an elementary and preliminary form of this reviewing process, explaining away ambiguities in the representation of the object trajectory, that will evidently necessitate further processing downstream the visual system. For instance, what we documented here could explain the ability of our visual system to detect objects based solely on the coherence of their spatio-temporal trajectory. In their seminal work, Watamaniuk and collaborators (1995) indeed showed that a single dot following a temporally coherent trajectory can be detected against a background of dots following a random walk, the only difference between signal and noise dots movement being their spatio-temporal coherence (Watamaniuk, McKee, and Grzywacz 1995). Computational studies suggested that this ability to detect coherent trajectories necessitates propagation of information in retinotopic reference frames (Perrinet and Masson 2012), in full accordance with our results.

### Local vs Global motion processing

The processing that we describe here clearly departs from classical motion integration documented in short-range apparent motion using random-dot kinetogram (Mikami, Newsome, and Wurtz 1986b, [a] 1986) In these stimuli, motion occurs and is evenly distributed within a stationary aperture typically covering a receptive field, and motion is extracted locally through motion energy detectors (Majaj, Carandini, and Movshon 2007; Pack et al. 2006). Simple L-NL hierarchical models account very well for the selective properties of neurons in V1 and MT in response to such kind of drifting or RDK stimuli (Rust et al. 2006; Carandini et al. 2005). However, there should be intrinsic differences in the processes involved in integrating local drifting motion vs global trajectory motion of a single object. Indeed, Hedges and collaborators (2011) have showed that MT receptive fields are only sensitive to local motion presented within stationary aperture, totally independent of the direction of long-range trajectory simulation in which these local motion stimuli are embedded (Hedges et al. 2011). We have very limited understanding of the processing actually required to extract motion information along a trajectory. The experiments of Watamaniuk and colleagues show that this processing cannot be simply integrated from large receptive field of downstream areas (Watamaniuk, McKee, and Grzywacz 1995). Here we suggest that the visual system can simply encodes the trajectory at mesoscopic level within retinotopic map.

### Encoding the motion trajectory in the retinotopic map for optimal read-out

The suppressive wave we documented decreases the residual activity evoked by the first stimulus, hereby shaping the dynamic response within the retinotopic map of V1 that could be read out as motion information by a downstream area. V4 or MT neurons have receptive fields whose retinotopic size encompasses the cortical region we imaged in this study. As shown by our read-out analysis (Fig. 8), those neurons will be able to simply detect this population-encoded direction selective motion information through motion energy detectors (Adelson and Bergen 1985). This signifies that V1 intra-cortical interactions would preformat the population representation of long-range apparent motion for an optimal read-out by downstream areas (Adelson and Bergen 1985; Mumford 1991, 1992). One intriguing consequence is that encoding of motion signal at the level of the population could be operated without specific extraction of motion signal at the level of local V1 neuronal receptive fields. Indeed, neurons with non-optimal direction preference or no direction selectivity could still participate into this population response by small variations of their response that would occur at the right moment depending on their position in the retinotopic space. In other words, V1 would have the possibility to encode multiple motion signals in parallel at local and global level. These results are in accordance with human fMRI experiments that showed that V1 is actively involved in the network that processes and represents the perceived illusory lrAM (Muckli et al. 2005).

### lrAM along ventral and dorsal streams, feedback vs horizontal propagation

In the visual cortex of the ferret, it was shown using VSDI, that lrAM induces feedback propagation of differential activity from area 21 down to area 17 (Roland et al. 2006). Similarly, using stimuli that could span a much large visual scale (16.5° spatial separation) and systematically larger cortical separations, it was suggested that human MT complex feedbacks on early visual cortices to process long-range apparent motion (Wibral et al. 2009; Vetter, Grosbras, and Muckli 2015). Areas on the ventral stream (LOC) seems to be also implicated in processing such stimuli (Zhuo et al. 2003). Ventral stream areas may actually be well suited since they will process the information about object through strong feedback interactions with V1 (Poort et al. 2012) and are as well strongly involved in motion processing (Roe et al. 2012; Ferrera, Rudolph, and Maunsell 1994). The experiment from (Hedges et al. 2011) indeed suggested that MT may not be the most appropriate area, at least in non-human primates, for extracting motion along a lrAM trajectory. It is important to consider though that, in all these studies, there are important difference in the spatial and a temporal scales of the lrAM has been presented that may affect the relative weight of intra-cortical and feedback mechanisms processing this information between and within the different visual areas (see Discussion in Reynaud, Masson, and Chavane 2012).

## Conclusion

As recently proposed by Muller et al. (2018), traveling waves within and between cortical areas can provide an advantageous framework for dynamic computations that will influence neuronal processing. However, in this review, it was also noted that clear functional roles of these waves have yet to be discovered. Here we show that two discrete stimuli generating the long-range apparent motion illusion, will induce multiple wave interactions resulting in propagation of suppression in a direction opposite to that of the AM stimulation. This suppression shapes the stimulus response and allows a decoder to keep track of the stimulus position along the motion trajectory. We believe that our work has revealed a first elementary step in how the brain links visual stimuli in space and time. Further work is needed to understand which areas, if any, is reading-out the population representation of motion trajectory on V1 retinotopic map and the relative role of intra- and inter-cortical interactions.

## Acknowledgments

The authors are thankful to Guillaume Masson, Andrew Meso, Eero Simoncelli, Dirk Jancke, Yves Frégnac, Cyril Monier, Lyle Muller and Tony Movshon for fruitful discussions during different phases of this work. They are also grateful to Marc Martin, Frédéric Barthélemy, Ivan Balansard and Luc Renaud for their assistance regarding experiments.

The authors acknowledge funding from the European Community (FET grants FACETS FP6-015879 and BrainScaleS FP7-269921), from la Fondation de l’œil (IA) and from the French National Research Agency (ANR Trajectory, ANR-15-CE37-0011-01, and ANR Horizontal V1, ANR-17-CE37-0006-02).

## Author Contributions

F.C. designed the research and supervised the project. S.C. and A.R. performed the experiments and S.C. analyzed the data. M.D. and Y.Z. designed and implemented the computational model in interaction with S.C. and F.C., while supervised by A.D.. F.C. and L.P. designed the decoding model and S.C. implemented it. S.C. prepared the figures and F.C. wrote the main manuscript text. All authors reviewed the manuscript.

## Materials and Methods

The experiments were conducted on two male rhesus macaque monkeys (macaca mulatta, aged 14 and 11 years old respectively for monkey WA and monkey BR) over a period of three years. The experimental protocols had been previously approved by the local Ethical Committee for Animal Research (approval A10/01/13, official national registration 71-French Ministry of Research) and all procedures complied with the French and European regulations for Animal Research as well as the Guidelines from the Society for Neuroscience.

### Surgical preparation and VSDI protocol

The monkeys were chronically implanted with a head-holder and a recording chamber located above the V1 and V2 cortical areas of the right hemisphere. After full recovery, the monkeys were trained to perform foveal fixation of a small red target presented over different static and moving backgrounds for up to 2-3s, with their head fixed. Once a good fixation behavior was achieved, a third surgery was performed. The dura was removed surgically over the recording aperture (18mm diameter) and a silicon-made artificial dura was inserted under aseptic conditions to allow for a good optical access to the cortex over the whole period of weekly recordings. Before each recording session conducted in awake animal, the cortical surface was stained with the Voltage Sensitive Dye (VSD) RH-1691 (Optical Imaging ©) with the following procedure: The optical chamber was open, artificial dura-mater was removed and cortical surface was cleaned under strict sterile conditions. The dye solution was prepared in artificial cerebrospinal fluid (aCSF) at a concentration of 0.2 mg/ml, and filtered through a 0.2μm filter. The recording chamber was filled with this solution and closed for three hours, corresponding to the time lapse needed for a correct cortical staining. The chamber was then rinsed thoroughly with filtered aCSF to remove any supernatant dye. Before imaging, the artificial dura was placed back in position and the chamber was closed with transparent agar and cover glass. Experimental control, data collection and eye position monitoring were performed by the ReX software (NEI-NIH) running under the QNX operating system (Hays et al., 1982). During each trial, the cortex was illuminated at 630 nm using epi-illumination and we recorded optical signals high-pass filtered at 665 nm during 999ms with a Dalstar camera (512×512 pixels resolution, frame rate of 110 Hz) driven by the Imager 3001 system (Optical Imaging ©). The beginning of both online behavioral control and image acquisition were heartbeat-triggered. The surgical preparation and VSD imaging protocol have been described elsewhere (Reynaud, Masson, and Chavane 2012; Muller et al. 2014).

### Behavioral task and visual stimulation

Monkeys were trained for a simple fixation task. For each experimental trial, the monkeys were required to fixate a central red dot within a precision window of 1°×1°. When correct fixation was achieved, the next heartbeat, detected with a pulse oximeter (Nonin 8600V), triggered the beginning of the acquisition window. A visual stimulus appeared 100 ms after this trigger after which a blank screen was presented, ending the trial. Each trial ran for 700 ms. If the monkey had maintained fixation up to the end of the acquisition period, a reward (fruit compote drop) was given. Otherwise, the trial was canceled, an alert sound was delivered and the procedure was re-initiated. The visual stimuli were computed on-line using VSG2/5 libraries and were displayed on a 22” CRT monitor at a resolution of 1024×768 pixels. Refresh rate was set to 100Hz. Viewing distance was of 57cm. Luminance values were linearized by mean of a look-up table. We used Gaussian blobs with standard deviation (controlling the spatial width) of 0.5°. They were presented at different positions, located at 0.5° or 2° on the left of the vertical meridian respectively for monkey WA and monkey BR, and between 1.5° and 4.5° below the horizontal meridian. We used different stimulus durations, 10 ms(1 frame), 50ms or 100ms and different interstimulus intervals (ISI) for the two-stroke apparent motion stimulations (from 20 to 100 ms). All stimuli (single blobs of different durations, lrAM sequences and two blank conditions i.e. where no visual stimulus) were randomly interleaved with an inter-trial interval of 8 seconds for dye bleaching prevention.

### Data analysis

Stacks of images were stored on hard-drives for offline analysis. The analysis was carried on with Matlab R2014a (The MathWorks Inc. ©) using the Optimization, Statistics and Signal Processing Toolboxes. VSD evoked responses to each stimulus were computed in three successive basic steps. First, the recorded value at each pixel was divided by the average value before stimulus onset (“frame 0 division”) to remove slow stimulus-independent fluctuations in illumination and background fluorescence levels. Second, this value was subsequently subtracted by the value obtained for the blank condition (“blank subtraction”) to eliminate most of the noise due to heartbeat and respiration. Third a linear detrending of the time series was applied to remove residual slow drifts induced by dye bleaching.

### Spatio-temporal representation (ST data)

For each time frame, activity was averaged across the x-dimension within the apparent-motion trajectory (e.g. *dotted rectangle at frame 216 ms* in Fig. 1, C-G) to provide a unique spatial cortical dimension as a function of time.

### Latency estimation

Response latency was defined as the point in time at which the signal derivative crossed a threshold set a 2.57 times (99% confidence) the SD of its baseline computed during a 100-ms-long window right before stimulus onset.

### Speed estimation

Within the ST representation, the speed of activity propagation was estimated by computing the slope of the linear regression between each latency estimate as a function of the cortical distance in the ST representation

### Data Fitting

For extracting the space and time constants of the VSD responses, we fitted the ST data in space (for each time frame) to a Gaussian function of the form:

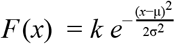

where σ, *k* and μ respectively denote the width (as the standard deviation), the amplitude and the spatial position of the Gaussian. We use the slope of the linear regression of μ(*t*) for quantifying the displacement of the response peak (see Fig. 4E).

In time (for each spatial point), the data was fitted to the combination of two halve Gaussian functions:

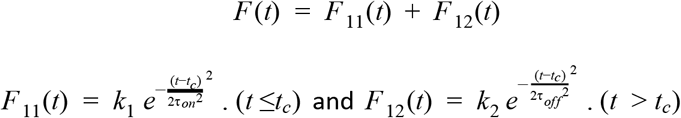

where τ_*on*_ and τ_*off*_ denote the time-constants of each half Gaussian, while *k*_1_, *k*_2_ and *t*_*c*_ are respectively their peak to peak amplitudes and the time of their common center.

### Statistical Procedure

We used a two-sample t-test procedure to test whether or not the distributions of the VSD response properties (i.e. space-constant, time-constants, latencies and cortical speed) were independent of stimulus duration or lrAM speed. p<0.01 is considered significant.

### Mean-field computational model

We consider a spatially extended ring model where every node of the ring represents the network activity of a large population of excitatory regular spiking (RS) and inhibitory fast spiking (FS) cells (see Fig. 5A). We consider Adaptive Exponential integrate and fire (AdExp) neurons evolving according to the following differential equations:

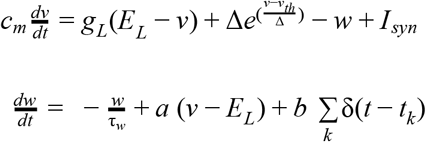

where *c*_*m*_ = 100 pF is the membrane capacity, v is the voltage of the neuron and, whenever *v* > *v*_*th*_ =− 50 mV at times *t*_k_, v is reset to its resting value *v*_rest_ =− 50 mV. The leak term has a conductance *g*_L_ = 10 nS and a reversal potential *E*_L_ =− 65 mV. The exponential term has a different strength for RS and FS cells, i.e. ∆ = 2 mV (∆ = 0.5 mV) for excitatory (inhibitory) cells. Inhibitory neurons do not have adaptation (a=b=0) while excitatory neurons have an adaptive dynamics with *a* = 4 nS, b=40 nS and τ_*w*_ = 500 ms. The synaptic current can be expressed as:

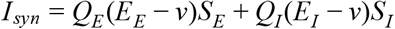

where 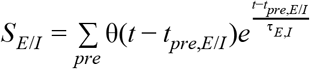 is the postsynaptic current due to all presynaptic excitatory/Inhibitory neurons spiking at time *t*_*pre*,*e*/*I*_ and *θ* is the Heaviside function. The reversal potentials are *E*_E_ = 0 mV and *E*_I_ =− 80 mV, the synaptic decays are equal for excitatory and inhibitory cells, τ _*E*,*I*_ = 5 ms. The quantal conductances are *Q*_E_ = 1 nS and *Q*_I_ = 5 nS. We then consider a random network with p=5% of connectivity and 80% of excitatory neurons.

The activity of the network is simulated using a mean field model, shown capable of quantitatively predicting the stationary activity of the network and its response to an external stimuli (Zerlaut et al. 2018). All together, the dynamical equations for the spatially extended ring model read:

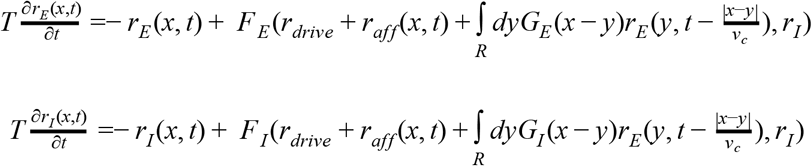

where *r*_*E*/*I*_ (*x*, *t*) is the population rate of excitatory/Inhibitory cells at the space-time position (x,t), *r*_aff_ (*x*, *t*) is the excitatory afferent input targeting both excitatory and inhibitory populations and *G*_*E*/*I*_ is the spatial connectivity in between subpopulations that we chose as Gaussian of width *l*_*exc*_ = 5 mm (excitation) and *l*_*inh*_ = 2.5 mm (inhibition). Moreover, *v*_*c*_ = 300 mm/s is the axonal conduction speed, *r*_drive_ an external time/space constant external drive and T=5ms is the decay time of population rate. The functions *F*_*E*,*I*_ are the transfer functions of excitatory/inhibitory neurons and are calculated according to a semi-analytical tool as in (Zerlaut et al. 2018) through an expansion in function of the three statistics of neurons voltage, i.e. its average μ*V*, its standard deviation σ*V* and its autocorrelation time τ *V*:

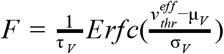

where *Erfc* is the error function and the effective threshold 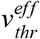 is expressed as a first order expansion with some fitting coefficients in function of (μ_*V*_, σ_*V*_, τ_*V*_). More details on this procedure can be found in Zerlaut et al. (Zerlaut et al. 2018). The values (μ_*V*_, σ_*V*_, τ_*V*_) are calculated from shot-noise theory (Daley and Vere-Jones 2007). Introducing the following quantities:

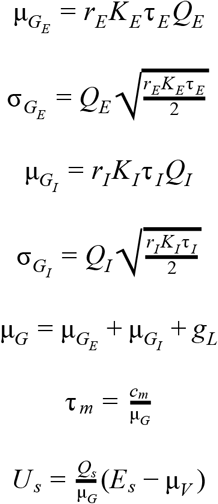

where *K*_*e*/*I*_ is the amount synapses related to pre-synaptic excitatory/inhibitory neurons (we consider a network of N=10000 neurons inside each node of the ring), we obtain the following equations for the voltage moments:

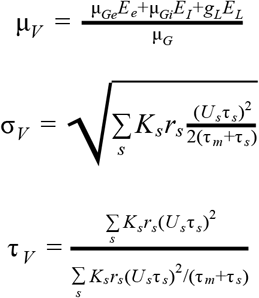

The afferent input has the following form:

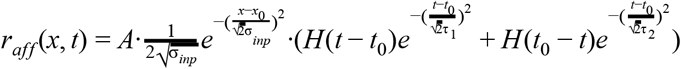

where A is the input amplitude, (*x*_0_, *t*_0_) the stimulus location. And H the heaviside function. The spatial extension of the stimuli is σ_*inp*_ = 3.5 mm, the time rise τ _1_ = 15 ms and the decay time τ _2_ = 90 ms. The time delay in between stimulus 1 and stimulus 2 is ∆_*t*_ = 100 ms (if not stated differently) and the spatial distance ∆_*x*_ = 7 mm. The VSDI signal is calculated as follows:

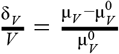

where 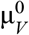 is the average voltage pre-stimuli.

#### CUBA model

The current based model is obtained by considering the following synaptic coupling:

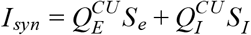

where 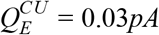 and 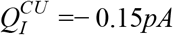 are the coupling with excitatory and inhibitory neurons. The rest of the parameters are the same. The voltage of the neurons is calculated accordingly, i.e.

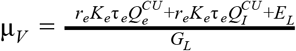

Also in this case we use the same methodology to estimate the neurons transfer function as done for the COBA model.

#### Different FS gain

In order to modify the gain of FS cells we manually change the transfer function *F*_*I*_(*r*_*E*_, *r*_*I*_). In practice, for any *r*_*I*_ we calculate the value 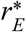 for which TF changes convexity. This gives us the slope 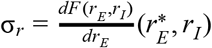 and the maximal value *F*_*max*_ that we estimate calculating F for very high rates (typically *r*_*E*_ = 200*Hz*). We then use the following function:

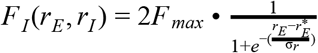

where we recall that 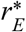 and ν_*r*_ change in function of *r*_*I*_. This permits us to have a sigmoidal form of the transfer function *F*. In order to change its slope we use a factor γ that scales the slope which becomes then γσ_*r*_. In Fig. 4 we use γ equal to 1.2 or 0.8.

### Decoding Model

The algorithm for the decoding model used in Figures 6 and 7 is detailed here. First, the ST data (i.e. space-time matrix) were whitened (i.e. spatially decorrelated and scaled) by applying a ZCA transformation. The whitening matrix was computed from the eigen-decomposition of the covariance matrix of the blank data. Next, the four spatial profiles (blank, stimulus 1, stimulus 2 and joint stimulus 1 and 2) were computed by averaging the corresponding ST response in a 50 ms-window around the time of maximum response and then normalized. The decoding of any ST data (e.g. the observed activity evoked by a 6.6 °/s two stroke apparent motion stimulus “*obs*” or its linearly predicted pattern “*pred*”) thus consisted in evaluating the likelihood that the spatial profile observed at one point in time of the data *A*(*x*, *t*) was best correlated with one of the four spatial profiles *S*_*j*_ with *j*∈{1 : 4}). This comes down to calculating the four probability *P*_*j*_(*t*) of the form:

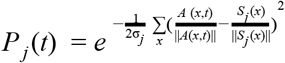

where σ_*j*_ is the averaged standard deviation of the residual activity between *A*(*x*, *t*) and *S*_*j*_ (*x*).

Then, we defined the explaining away index as the probability of detecting joint S1&S2 in the observed 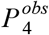 or 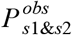 minus the probability of detecting joint S1&S2 in the linear prediction 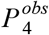 or 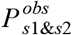 as follows:

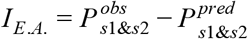

### Opponent motion energy model

To extract motion information from the population responses, we used the opponent motion energy model developed by (Adelson and Bergen 1985). Briefly, this model consists of combining quadrature pairs of spatial and temporal filters to obtain oriented spatio-temporal filters (i.e. Gabors) tuned in spatial frequency. The ranges of spatial and temporal frequencies were chosen so that the speed (i.e. FT/FS) of the resulting ST filters varies from 2 to 70 °/s and the scale (i.e. 1/FS) from 0.2 to 6 mm. It resulted in 64 (FS,FT) couples representing 8 different speeds and scales. For each couple, we obtained two filters tuned for upward motion and two filters tuned for downward motion. The outputs of quadrature pairs of such filters are then squared and summed to give a phase-independent measure of local motion energy for both directions (i.e MEu and MEd values). Lastly, the opponent motion stage computes the difference between the oriented opposite energies (i.e. OME values). Note that before applying the OME model, the ST data were first normalized and passed through a non-linearity to account for the VSD to spike rate transformation as proposed by (Chen, Palmer, and Seidemann 2012):

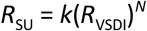

where *R*_SU_ and *R*_VSDI_ are respectively the average firing rate and the average normalized VSDI response, *k* is a constant and *N* is an exponent. Here we took *k* = 10 and *N* = 3.8.

Finally, for each ST position on the map, we could extract the velocity of the filter that generated the strongest OME and provide a ST velocity map representation (Fig. 8C-D) with velocity and amplitude as color hue and color intensity respectively. We then averaged the optimal velocity within a ST region of interest, spatially between S1 and S2’s center positions and in time from 10 to 200 ms after stimulus 2 onset, to report a single value of filter speed for each AM speed condition (Fig. 8D-E). The direction-selectivity index is given by:

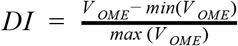

where*V*_*OME*_ is to the amplitude of the OME.

